# Diagnostic outcomes of exome gene panel sequencing in patients with unusual syndromic cleft lip/palate phenotypes

**DOI:** 10.1101/465179

**Authors:** M. Reza Jabalamelil

## Abstract

Orofacial clefting (OFC) is a common craniofacial birth defect that has a prevalence of 1.2 in 1,000 live births. Syndromic OFC in which patients present with additional developmental deficits are identified to have a strong genetic component. We applied exome gene panel sequencing in a cohort of 14 Colombian patients identified at Operation Smile in Bogota, Colombia, with syndromic orofacial clefting phenotypes and additional features initially suggesting Aarskog-Scott syndrome (AAS). Gene panel sequencing failed to identify any causal variants in the *FGD1* gene which underlies this condition, but variants in a number of genes suggest alternative clinical diagnoses across five patients (~ 36%). The novel variants identified here maps to developmentally important genes including *SRCAP, OFD1, NIPBL, GRIN2A* and *KMT2D6*. Our result demonstrates the extensive heterogeneity underlying OFC and emphasises the need for systematic phenotyping of patients with rare conditions. Gene panel sequencing has the potential to cost-effectively resolve ambiguous clinical diagnoses, but rigorous attention should be paid to gene coverage as our results suggest highly variable read depths across individual gene exons and this may reduce the quality of variant calls at specific locations.

## Introduction

Orofacial clefting (OFC) is a common congenital abnormality which has a prevalence of between one and two individuals per thousand live births [1]. Clefting arises from incomplete fusion of primordial plates involved in orofacial development. Cases can be classified in terms of whether they have cleft lip with or without cleft palate (CL/P), or cleft palate only (CPO). However, the distinction between nonsyndromic (isolated) forms and syndromic conditions (with additional anomalies present) is particularly important. Nonsyndromic OFC is a complex phenotype arising through interaction between multiple genes and environmental factors. In contrast, syndromic conditions, of which about 400 have been described [1], are generally single gene disorders which are therefore amenable to analysis of case samples using next-generation sequencing (NGS).

The study by Arias Uruena *et al*. [2] describes the clinical features of patients identified at the Operation Smile clinic in Bogota, Colombia. The patient cohort includes individuals with both non-syndromic and syndromic forms of OFC. Analysis of 311 patients attending the clinic over the period April 2012-July 2013 identified 59 patients with syndromic conditions which were provisionally classified according to their clinical presentation. An unexpected finding was the observation that 14 of these patients (24%) showed a number of phenotypes suggesting a possible diagnosis of Aarskog-Scott syndrome (AAS, OMIM #305400). This patient group had the highest frequency amongst 27 syndromes recorded, matched by Velocardiofacial syndrome (n=10), with smaller numbers of other syndromes including Pierre Robin (n=5), Van der Woude 1 (n=3) and Orofaciodigital 1 (n=3) and Ectodermal Dysplasia and Hypohidrotic 1 (n=3). In contrast, previous studies found that Van der Woude syndrome (VWS1, #119300) is the most frequent syndromic condition amongst OFC patients [3–5]. It was suggested that local geographical and ethnic factors in this Colombian population might account for the atypical distribution of syndromic forms identified.

Aarskog-Scott syndrome (also known as faciogenital dysplasia) is a complex developmental disorder with diagnosis normally established through the Teebi criteria [6] for which short stature, hypertelorism and fold of the lower lip are features of most cases [7]. Brachydactyly, interdigital webbing, shawl scrotum, long philtrum and mild facial hypoplasia are secondary features observed in ~80% of cases. Additional phenotypic manifestations including cryptorchidism, inguinal hernia, downward eye slant and ptosis are present in a fraction of patients [8]. AAS patients usually present with delayed growth in early childhood but achieve developmental milestones later in life [9]. AAS predominantly affects males with the phenotypic complications being attenuated in females. Established genetic variants for AAS are located on the proximal short arm of chromosome X (Xq11.22) and comprise at least 61 mutations spanning 18 exons of the *FGD1* (FYVE, RhoGEF and PH domain containing 1) gene [8]. Patients with proven FGD1 mutations have a prevalence of approximately 1/25000 in the general population [10]. Despite this, cases with confirmed *FGD1* mutations only comprise ~20% of cases with reported AAS phenotypes [11]. This substantially incomplete understanding of the genetic basis of the condition may reflect genetic heterogeneity and/or phenotypic heterogeneity with a degree of sharing of phenotypic features between several conditions. For cases in the Colombian sample, the predominant OFC phenotype is atypical for a diagnosis of AAS, and we, therefore, undertook gene panel sequencing of 14 of the cases intending to establishing molecular diagnoses. We also evaluated the utility of gene-panel sequencing for resolving molecular diagnoses in general and consider patterns of sequence coverage across the *FGD1* gene in particular as an illustrative example.

## Materials and methods

### 0.1 Patients phenotype

DNA samples from 14 unrelated male patients each with a tentative clinical diagnosis of AAS were obtained through the Operation Smile clinic in Bogota, Colombia (Table 1). All patients presented with either unilateral or bilateral cleft lip (CL) and nine patients also presented with complete cleft palate (CP) and one with incomplete CP. Shawl Scrotum (SC) was noted for 13 patients and cryptorchidism in four patients. The majority of patients showed evidence of orbital hypertelorism. Some patients showed reduced height and/or weight for their developmental stage, and nine patients were recognised as having the developmental delay. Details of these and other phenotypes are presented in Table 1.

**Table 1.** Phenotypic features of the 14 patients with primary diagnosis of AAS considered for exome gene-pane1 sequencing.

| ID | Age | Cleft Lip | Cleft Palate | Shawl Scrotum | Cryptorchidism | Telecanthus | Hypertelorism | Short Stature | Developmental Delay | Additional Features |
| --- | --- | --- | --- | --- | --- | --- | --- | --- | --- | --- |
| CL021 | 11 yrs | Bilateral | Incomplete | + | - | - | + | + | + |  |
| CL022 | 13 mths | Bilateral | Complete | + | + | + | + | + | - |  |
| CL024 | ? | Bilateral | - | - | - | + | + | - | + |  |
| CL025 | 3 yrs | Unilateral | - | + | + | + | + | + | + | Anophthalmia,<br>Esophageal atresia |
| CL027 | 9 yrs | Unilateral | Complete | + | - | - | + | - | - |  |
| CL028 | ? | Unilateral | Complete | + | - | - | - | + | + |  |
| CL030 | 2 mths | Unilateral | Complete | + | - | + | + | - | - |  |
| CL033 | 10 yrs | Unilateral | Complete | + | - | + | + | - | - | Clinodactyly |
| CL035 | 8 mths | Unilateral | Complete | + | - | - | - | - | + |  |
| CL036 | 5 yrs | Unilateral | Complete | + | + | + | + | + | + | Clinodactyly,<br>Heart murmur |
| CL037 | ? | Unilateral | Complete | + | - | + | + | - | - | Left club foot |
| CL038 | ? | Unilateral | Complete | + | - | + | + | - | + |  |
| CL039 | 8 yrs | Unilateral | Complete | + | - | - | + | - | + |  |
| CL040 | 36 yrs | Bilateral | Complete | + | + | - | - | + | + | Lip melanosis |

### 0.2 Gene panel sequencing

Targeted exome enrichment was carried out using the TruSight One sequencing panel (Illumina, San Diego, CA, USA). The panel covers exonic regions of 4,813 disease-causing genes associated with clinical phenotypes that are indexed in the HGMD database [12] and OMIM [13]. Library preparation was carried out using the Nextera Rapid Capture Enrichment protocol (Illumina, San Diego, CA, USA). In brief, 50 ng of genomic DNA from each sample was enzymatically fragmented and tagged with adaptor sequences. Tagmented amplicons were then dually indexed and amplified through 10 cycles of PCR reactions. Cleaned libraries were hybridised to biotinylated probes specific to target regions and enriched by streptavidin beads. Sequencing-ready fragments were then magnetically pooled and eluted from beads to give a mean fragment size of ~300 bp. Sequencing was performed at the Wessex Investigational Sciences Hub laboratory (WISH lab, University of Southampton, UK) on a HiSeq2000 platform (Illumina, San Diego, CA, USA).

### 0.3 Sequence data processing and quality control

The quality of raw sequence data was checked, and high-quality reads were aligned to the human reference genome GRCH37 (hg19) using BWA aligner (v0.7.12). Following alignment, duplicate reads were marked and discarded using Picard tools (v1.97). The base quality scores were recalibrated by GATK (v3.6) BaseRecaliberator module and local realignment around indels was carried out using GATK IndelRealigner. In order to verify that samples are not contaminated, the Binary Alignment Map (BAM) files were checked by VerifyBAMID (v.1.1.3), and FREEMIX contamination score was used to confirm lack of contamination in samples. Variant calling for each sample was contemporaneously carried out in Samtools [14] v1.3.2 and GATK (v3.6) HaplotypeCaller [15]. The resultant VCF files from the two variant callers from each sample were then merged and annotated using ANNOVAR [16] (2015Dec14 release). Variant sites were defined according to the GRCh37/hg19 genome build, and known polymorphisms were annotated according to dbSNP [17] build 139. The 1,000 Genomes project phase 3 dataset [18] along with NHLBI Exome Sequencing Project (ESP) and Exome Aggregation Consortium (ExAC) datasets [19] were used for filter-based annotation. Furthermore, the in-house exome database was also queried to incorporate allele frequencies (AF) from ~ 460 regional WES analysed patients unaffected by CLP. Human RefSeq transcript dataset (GRCh37.p10) was used for gene-based annotation and pathogenicity scores and conservation scores for variant sites were computed using prediction algorithms including M-CAP [20], PolyPhen2 [21], SIFT [22], LRT [23], FATHMM [24], RadialSVM [25], CADD [26], Phylop [27] and GERP^++^ [28]. For variants outside the exon boundaries, the Ensemble Variant Effect Predictor (VEP) [29] MaxEnt plugin was used to quantify the impact of the mutational event on the splicing process. Splicing variants that achieved the MaxEnt significance threshold was further evaluated in the Human Splicing Finder (HSF, v.2.4.1) [30]. Ultimately, the HGMD record for disease-causing genes from the HGMD v.2016.2 VCF file was also added to the variant annotation.

Out of the 14 samples considered for variant analysis, one exome sample was discarded early on due to the discrepancy between the inferred gender from X-chromosome heterozygosity and the reported gender from the clinic. As for the remaining 13 samples, only eight exome samples fulfilled the QC criteria, and variant analysis was carried out for these samples (Table 2). All the QC-passed samples achieved >93% coverage of target sequences with a read depth of at least 20X and >57% of sequence reads mapped to TruSight One targets for all samples. The number of called DNA variants was reasonably consistent across samples in the range 11.4K to 12.9K for each sample. Contamination rates for all samples were consistently well below the conventional 2% threshold. The level of heterozygosity on the X chromosome was used as a check for sample gender, and the low percentage of heterozygous variants (<31% for all samples) mapping to the X chromosome confirmed sample gender as male, consistent with phenotypic data (Table 1).

**Table 2.** Quality Contro1 Measures for the eight sequenced patient samples

| Sample ID | Number of called<br>sequence variants | %X chromosome<br>Heterozygosity | Number<br>of reads | % Reads mapped to<br>TruSight One targets | % of target sequence<br>covered at 20X | verifyBamID f<br>reemix score % |
| --- | --- | --- | --- | --- | --- | --- |
| CL021 | 12101 | 23.02 | 30010619 | 67.08 | 96.58 | 0.010 |
| CL022 | 12312 | 21.38 | 42409723 | 57.77 | 97.13 | 0.019 |
| CL025 | 11431 | 19.01 | 29788440 | 70.78 | 93.44 | 0.027 |
| CL027 | 12371 | 30.72 | 32060201 | 60.92 | 96.50 | 0.013 |
| CL033 | 12364 | 22.56 | 48216232 | 64.31 | 97.79 | 0.011 |
| CL035 | 12203 | 21.42 | 29357251 | 64.89 | 96.83 | 0.011 |
| CL036 | 12939 | 15.43 | 42758969 | 64.18 | 97.64 | 0.009 |
| CL039 | 12127 | 21.76 | 28421558 | 64.44 | 95.69 | 0.011 |

### 0.4 Analysis of sequence read-depth coverage of the *FGD1* gene

We evaluated patterns of sequence coverage across the *FGD1* to evaluate the consistency of coverage across individual exons as an illustrative example. We obtained batch normalised sequence read depth for each of the 18 exons of the *FGD1* gene in all samples with consistent gender assignment (n=13). From this, we produced a heatmap plot of read-depth per exon for all these samples. Secondly, we compared the read-depth by exon profiles to the coverage of the gene exons in an independent sample of six patients exome sequenced using the Agilent SureSelect Human All Exon Capture kit v 5.0 to establish whether the coverage profiles of the *FGD1* gene are specific to gene-panel sequencing or representative of other targeted sequencing panels. Thirdly, we compared normalised read-depth profiles by exon with an independent sample of 18 cases sequenced on the same panel. From this, we produced a box-plot analysis comparing the eight patient samples with the 18 ‘controls’. Finally, we examined the distribution of unique (independent) single nucleotide variants identified in the coding region of the *FGD1* gene to determine whether patterns of read coverage of individual exons might impact identification of causal variation. The data used were extracted from the LOVD 3.0 *FGD1* database.

### 0.5 Variant filtering and analysis

Gene panel sequencing yielded 11,431–12,939 variants per sample (Table 2). Given the diversity of phenotypic features in each patient, and the possibility that different syndromes are involved, filtering and variant analysis considered a wide range of potential genes as potentially causal for further analysis. We focused on genes previously implicated in OFC syndromes and any other genes associated with shawl scrotum and cryptorchidism phenotypes. We considered variants in a comprehensive set of 918 genes combined from three searches as follows: (1) A list of 112 candidate genes prepared by literature search, nucleating from HGMD v.2016.2 [12], using the following keywords (and Human Phenotype Ontology, HPO terms): Cleft, Shawl Scrotum (HP:0000049) and Cryptorchidism (HP:0000028); (2) A list of 20 additional genes implicated in Shawl Scrotum (HP:0000049) and Cryptorchidism (HP:0000028) phenotypes retrieved from the Harmonizome (Ma’ayan Laboratory of Computational Systems Biology); (3) A list of 787 additional genes prepared from the OMIM database (January 2017 update), ClinVar (January 2017 update), Orphanet and the DDDG2P database using a set of 11 HPO terms as follows: mild global developmental delay (HP:0011342), hyperactivity (HP:0000752), intellectual disability (HP:0001249), unilateral or bilateral cleft lip (HP:0100336, HP:0100333), complete or incomplete cleft palate (HP:0000175), telecanthus (HP:0000506), hypertelorism (HP:0000316), shawl scrotum (HP:0000049), cryptorchidism (HP:0000028) and clinodactyly (HP:0000028).

Variant annotation revealed that no variants with known clinical effects related to any of the phenotypes were present in the sample. With the expectation that the variants associated with these syndromic phenotypes are likely to be rare and highly penetrant, we further considered only novel variants. Additional variants were excluded using the following criteria: (1) All synonymous variants located outside of the exon-intron boundaries were excluded. Synonymous variants within 10bp of either donor-splice site or acceptor-splice were retained and analyzed as potential splicing variants; (2) Variants located beyond 10 base pairs 5′ or 3′ of exons were dealt with as noncoding variants and excluded; (3) Non-frameshift deletion/insertion variants were excluded as they were unlikely to be causal in these cases; (4) Variants with read depth of coverage of less than 10 were excluded from further consideration as potential false positive; (5) All novel variants were examined visually in IGV and any which suggested low-quality genotype calls and/or errors in alignment due to, for example, extensive homopolymer sequences in the region, were excluded from further consideration. We considered evidence of pathogenicity using four functional prediction models (SIFT, PP-2, LRT and FATHMM), two conservation models (GERP^++^ and Phylop) and three combined scores (M-CAP, CADD and RadialSVM).

To priorities candidate causal variants for each individual a score which combines evidence across the main predictive models was used to generate a ranked order (Table 3). For each variant the predictive scores from the selected models were transformed to the single combined score Ψ_*i*_ according to the formula:

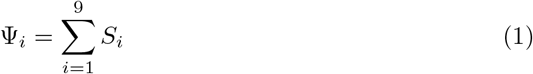

where *S_i_* is the score function defined as:

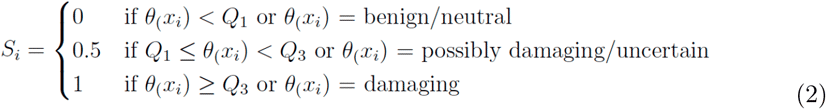

**Table 3.**
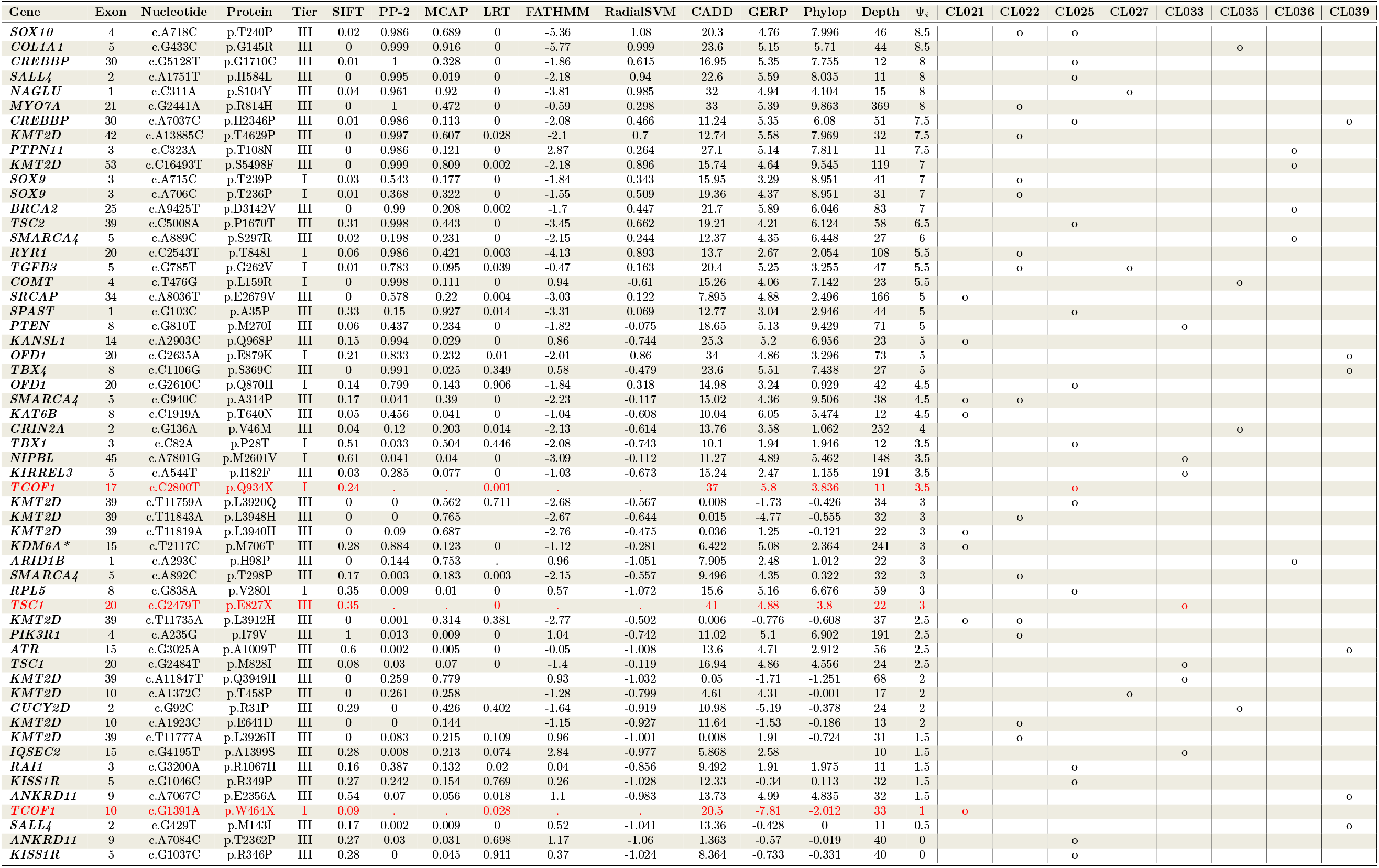
Novel variants with the dominant pattern of inheritance identified across the tiered analysis. Variants are ranked according to their

The *θ*(*x_i_*) is the pathogenicity or conservation score for variant x for the *i_th_* predictive model and Q denotes quartile range for scores from the M-CAP, CADD, GERP++ and Phylop models. Variants were then ordered from the largest (more pathogenic) to the smallest (less pathogenic). Considering that prediction scores for stop-gain mutations are not available from the Polyphen2, M-CAP, FATHMM and RadialSVM models, the potential pathogenicity of these variants was evaluated individually.

## Results

### 0.6 *FGD1* gene coverage

Figure 1 presents the heatmap of normalised read-depth coverage across all 18 exons of the *FGD1* gene for the samples. Also shown here is the corresponding data from whole exome sequences in six independent samples. These data suggest that the TruSight One capture kit does not provide a uniform capture across all 18 exons of this gene. Evidence from the independent samples (Figure 1.b) also suggests that coverage is far from uniform with generally close alignment between the two kits. In particular exons 5,6,7,10,14 and 16 are noticeably poorly covered.

**Figure 1.**
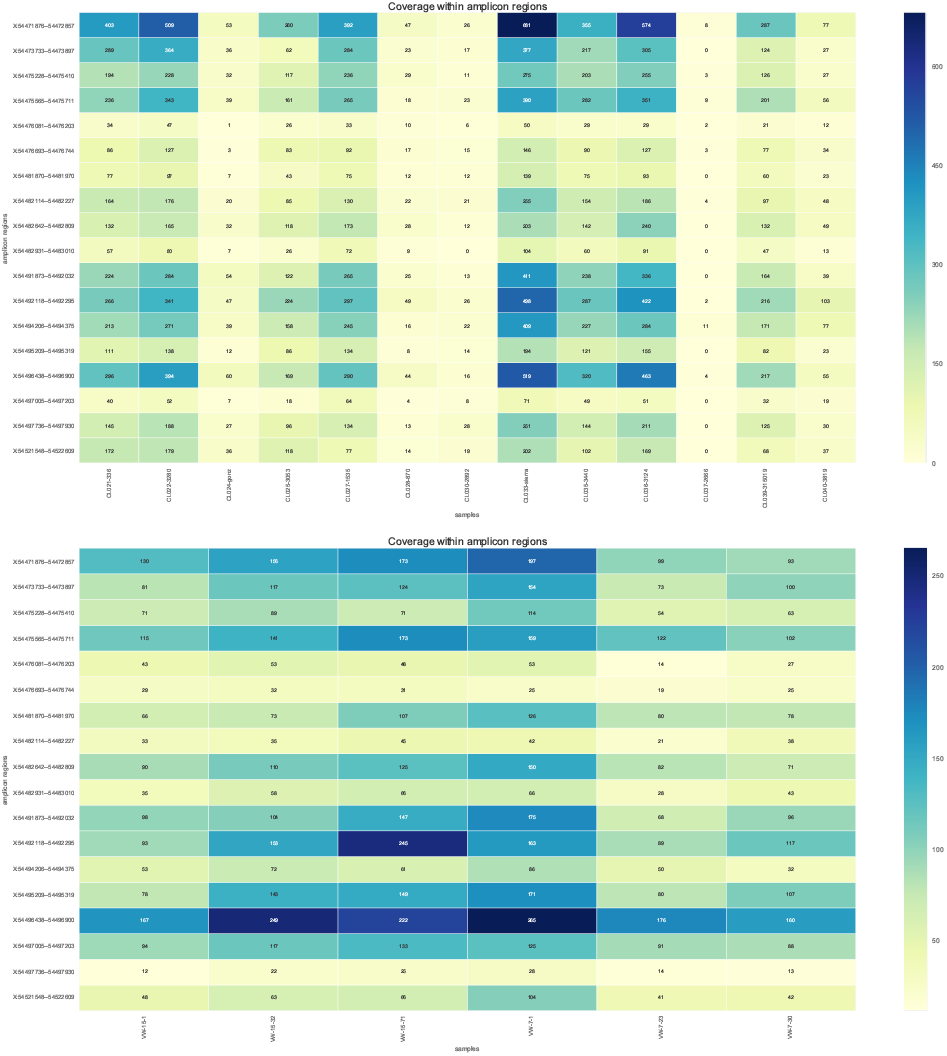
Heatmap plot of red depth per exon of FGD1 gene. a. The heatmap coverage plot of 13 OFD patients who underwent gene-panel sequencing; b. The heatmap coverage plot of six independent samples who were whole-exome sequenced using the Agilent SureSelect Human All Exon Capture kit v 5.0.

The box-plot (Figure 2) compares normalised read depths by exon for the eight samples with 18 control samples tested on the same gene panel. There is no significant difference between normalised read depths by exon for the two samples (*p*=0.99, one-tailed t-test) and similarly, uneven coverage is apparent.

**Figure 2.**
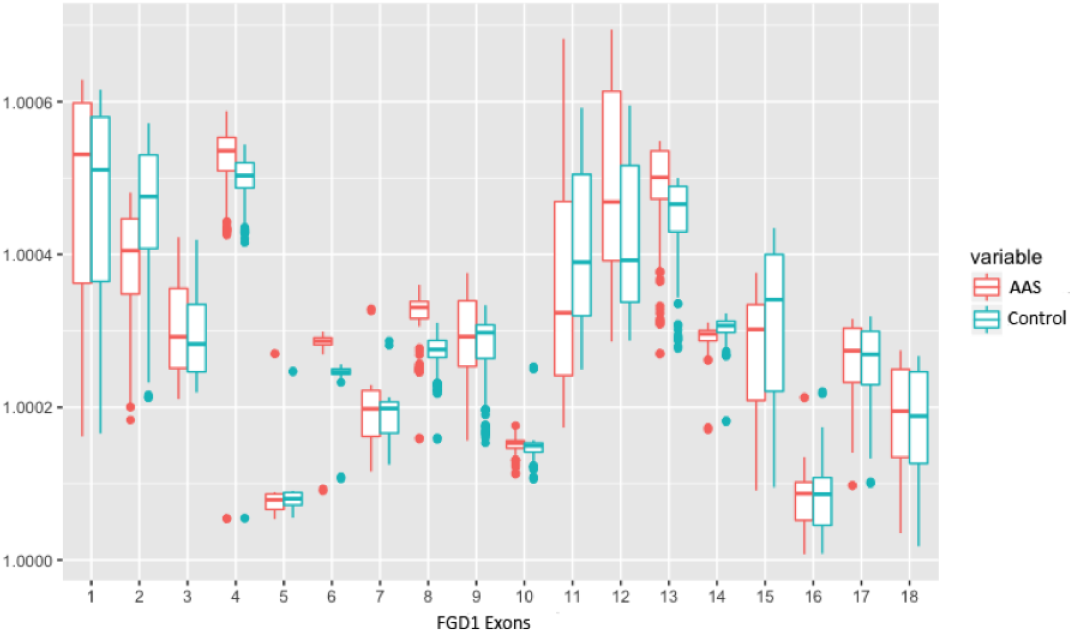
Normalised coverage across the *FGD1*in OFD patints (red) compared to 18 controls who were sequenced on the same panel (cyan

The results demonstrate that the uneven coverage pattern is neither capture kit nor sample specific and may reflect variation in the underlying sequence. We also considered the distribution of known unique AAS causal variants by *FGD1* gene exon (Figure 3). This shows that the highest number of unique causal SNVs map to exon 6 which is an exon with notable low read depth on both gene panel and exomes (Figures 1 and 2). These observations highlight the possibility that gene-panel or exome screening of samples for causal *FGD1* mutations may miss a causal variant.

**Figure 3.**
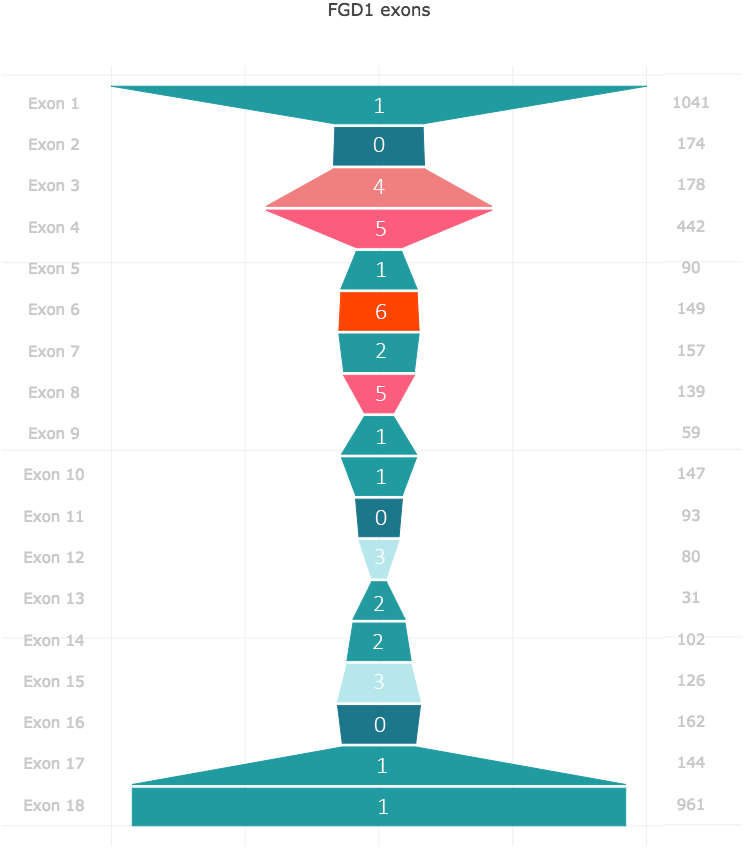
Frequency and localization of 38 novel SNVs identified in the coding region of *FGD1* gene. The vertical axis on the left identifies the exon number and the numbers on the right axis represent the size of each exon in bp. Trapezoids represent the relative size of exons and the numbers inside them show the total number of novel SNVs identified in each exon (Data extracted from the LOVD 3.0 FGD1 database)

### Variant Analysis

Table 3 lists 57 heterozygous variants across the eight cases Analysis by individual patient as follows:

#### Patient CL021

The patient is a 11 year male offspring of a non-consanguineous marriage presented with developmental delay (HP:0011342), incomplete cleft palate (HP:0000175), bilateral cleft lip (HP:0100336), hypertelorism (HP:0000316) and shawl scrotum (HP:0000049). He was delivered at 35 weeks of gestation and had low birth weight (~ 1100g) at the time of delivery. The proband’s parents are healthy, and CL feature is only present in his second-degree cousin. Eight variants were identified as possibly pathogenic in the proband *CL021* (Table 3).

The ranking strategy applied to prioritise putatively causal variants revealed the *SRCAP*:c.8036A>T as the top candidate. SRCAP is the activator of *CREBBP* and mutations of *CREBBP* have been identified to underlie Rubinstein-Taybi syndrome 1 (RSTS1, OMIM #180849). Mutations of the *SRCAP* have been described to underlie Floating-Harbor syndrome (FLHS, #136140) [31]. Patients with *SRCAP* mutation present with delayed bone age and speech development, short stature, triangular faces and deep-set eyes with long eyelashes. The ears in FLHS patients are posteriorly rotated, however the facial characteristics of the patients are age-related and change over the time. Unilateral cleft and cryptorchidism have also been reported in the context of the disease [31]. The majority of nonsense and frameshift *SRCAP* mutations that have been identified in FLHS patients map to the exon 34 of the gene [32]. As exon 34 is the last exon of the *SRCAP* gene, these mutations escape the nonsense-mediated mRNA decay and lead to pathogenicity [33]. Given that identified mutation maps to the exon 34 of *SRCAP* and the extent of phenotypic similarity between the proband’s phenotype and FLHS features, the functional impact of the variant merits further investigation.

Furthermore, hemizygous mutations of *KDM6A* (rank 6) have been identified to underlie X-linked dominant Kabuki syndrome 2 (KABUK2, OMIM # 300867). Patients with *KDM6A* mutation present with congenital mental retardation short stature, scoliosis and a range of facial dysmorphisms including long eyelashes, arched and sparse eyebrows, broad nasal bridge and cleft palate [34]. Miyake *et al*. demonstrated that *KDM6A* mutations underlie 6.2% of KABUK2 incidence. They also suggested that arched eyebrows are less common among *KDM6A* mutants but short stature and postnatal growth retardation is the consistent feature among all patients with *KDM6A* mutations. Genitourinary anomalies are common among KABUK2 patients but shawl scrotum has never been reported in the context of the disease. Furthermore, congenital mental retardation in combination with multisystemic anomalies involving heart and immune system is common among Kabuki patients. Given that the proband *CL021* does not present with intellectual disability and also considering the extent of phenotypic dissimilarity between the patient and KABUK2 syndrome, it is unlikely that the KDM6A:c.2117T>C variant is causal. This has been also reflected in the reduced combined pathogenicity score computed for this variant (Ψ_*i*_=3) and therefore this variant was excluded from further follow up.

The visual inspection of reads using IGV for the remaining variants including *KANSL1*: c.2903A>C (rank 2), *SMARCAĄ*: c.G940C (rank 3), *KAT6B*:c.1919C>A (rank 4), *KMT2D*:c.11735T>A (rank 5), *KMT2D*:c.11819T>A (rank 7) and *TCOF1*: c.1391G > A (rank 8) revealed low genotype quality calls and erroneous alignment. Based on this evidence, variants identified in these genes are probably spurious and therefore excluded from further follow up.

In conclusion, targeted exome sequencing and tiered filtering identified *SRCAP*:c.8036A>T as a putatively causal variant in the patient *CL021*.

#### Patient CL022

The patient is a 13 month old male offspring of a non-consanguineous marriage with normal development and no history of CLP in the family. Patient’s main features include short stature and low birth weight, bilateral cleft lip (HP:0100336), complete cleft palate (HP:0100336), hypertelorism (HP:0000316), telecanthus (HP:0000506), shawl scrotum (HP:0000049) and cryptorchidism (HP:0000028) as described in the Table 1. Fourteen variants were prioritised through the filtering strategy (Table 3). The visual inspection of reads in IGV to confirm the authenticity of variants revealed the top three candidates as MYO7A:c.2441G>A (rank 2), *RYR1*: c.2543C>T (rank 6) and *PIK3R1*: c.235A>G (rank 12).

*MYO7A* encodes an unconventional myosin with a very short tail that plays an important role in cellular movements [35]. Heterozygous mutations of *MYO7A* are identified to underlie nonsyndromic progressive hearing loss known as autosomal dominant deafness-11 (DFNA11, OMIM # 601317) [36]. Considering the lack of phenotypic similarity between the DFNA11 and the patient’s main features and also absence of vestibular symptoms in the proband *CL022*, it is highly unlikely that MYO7A:c.2441G>A is causal in the context of the disease. The variant, therefore, was excluded from further follow-up.

*RYR1* encodes a ryanodine receptor which acts as a calcium release channel in the sarcoplasmic reticulum of skeletal muscles [37]. Heterozygous mutations of *RYR1* have been identified to underlie Central core disease (CCD, OMIM # 117000) [38] and a form of malignant hyperthermia known as King-Denborough syndrome (OMIM # 145600) [39]. Facial and genitourinary dysmorphisms have never been reported in the context of CCD or King-Denborough syndrome. Furthermore, patients with pathological *RYR1* usually present with additional muscle and soft tissue involvement including neonatal hypotonia, muscle atrophy and muscle weakness in CCD or muscle rigidity and rhabdomyolysis in King-Denborough syndrome [40]. These phenotypes are absent in the proband *CL022* and therefore, given the inconsistency of symptoms between the patient’s main features and the symptoms associated with the *RYR1* mutations, it is unlikely that the *RYR1*: c.2543C>T is aetiological in the context of the disease.

Heterozygous mutations of *PIK3R1* are identified to underlie the autosomal dominant SHORT syndrome (OMIM # 269880). *PIK3R1* encodes the phosphatidylinositol 3-kinase that plays an important role in growth signalling pathways. This lipid kinase phosphorylates the inositol ring of phosphatidylinositol and thereby triggers the second messenger in the insulin signalling pathway [41]. Patients with the SHORT syndrome are present with short stature, triangular faces, micrognathia, telecanthus and deep-set eyes [42]. Bone maturation and teething are usually delayed among SHORT patients, but patients are intellectually normal [43]. CLP and genitourinary abnormalities including shawl scrotum and cryptorchidism have never been reported in the context of the SHORT syndrome. Despite the presence of telecanthus and short stature in the proband *CL022*, the phenotypic similarity between the SHORT syndrome and patients main features are negligible and therefore the *PIK3R1*: c.235A>G is unlikely to be relevant to the patient’s phenotype.

In conclusion, neither of patients phenotype was consistent with the syndromes associated to these genes and therefore a molecular diagnosis could not be established in this patient.

#### Patient CL025

The patient is a 3-year-old male child of a non-consanguineous marriage with a negative history of CLP in the family. The patient’s main features include developmental delay (HP:0011342), unilateral cleft lip (HP:0100333), hypertelorism (HP:0000316), telecanthus (HP:0000506), shawl scrotum (HP:0000049) and cryptorchidism (HP:0000028). The patient is also present with anophthalmia (HP:0000528) and oesophagal atresia (HP:0002032) (Table 1). Fifteen novel variants were prioritised through the filtering strategy, of which two variants including *CREBBP*:c.7037A>C (rank 4) and *OFD1*: c.2610G>C (rank 7) were identified as putatively causal with respect to the patient’s phenotype (Table 3).

Heterozygous mutations of *CREBBP* have been identified to underlie autosomal dominant Rubinstein-Taybi syndrome 1 (RSTS1, # 180849). Patients with pathogenic *CREBBP* mutations present with multiple congenital anomalies including dysmorphic facial features, broad and angulated thumbs, postnatal growth deficiency, shawl scrotum and cryptorchidism [44]. Facial dysmorphisms in RSTS1 patients Occasionally involves cleft lip [45], but anophthalmia has never been reported in the context of the disease. Given the lack of convincing phenotypic similarity between the patient’s main features and RSTS1, it seems unlikely that *CREBBP*:c.7037A>C underlies the patient’s condition.

Heterozygous mutations of *OFD1* have been identified to underlie the orofaciodigital syndrome I (OFD1, # 311200) with X-linked dominant pattern of inheritance. OFD1 patients are typically present with a range of malformations in the face, hand and oral cavity. Main features of the disease include short stature, cleft lip and cleft palate, hypertelorism, telecanthus and polycystic kidney disease [46,47]. Shawl scrotum and cryptorchidism have been rarely documented in the context of the disease as pathogenic *OFD1* mutations leads to premature death in hemizygous males [47]. Intriguingly, Tsurusaki *et al*. [48] reported a lethal case hemizygous male in a family ascertained for *OFD1* mutation in a hemizygous male with preterm delivery (33 weeks). The patient presented with cleft lip, soft cleft palate, hypertelorism, microphthalmia, micropenis and cryptorchidism. Tsurusaki *et al*. did not report esophageal atresia in the patient but reported that the patient died 11 hours after birth due to laryngeal anomalies. Genetic analysis of another male patient from the same family with similar phenotype and esophageal abnormalities who were also delivered preterm and died one day after birth revealed a pathogenic splicing mutation at *OFD1*: c.2388+1G>C [48]. Given extensive phenotypic similarity between the the patients described by Tsurusaki *et al*. and the proband CL025 main features it is possible that *OFD1*: c.2610G>C (Ψ_*i*_ =4.5) underlies the patient’s condition. One possible explanation for the lack of lethality in the proband *CL025* could be the mosaic nature of the mutation in the patient. In fact, the *OFD1*: c.2610G>C variant is supported by 8 reads out of a total read depth of 42 suggesting 19% mosaicism. Should mosaicism underlie the reduced penetrance in the patient CL025, *de novo* mutation of *OFD1* could be accounted for the patient’s phenotype.

#### Patient CL027

The patient is a 9-year-old boy of a consanguineous marriage where parents are first cousins and the patient’s third-degree cousin present with CLP. The patient is developmentally normal with his weight and height percentiles are in the normal range. The patient’s main features include unilateral cleft lip (HP:0100333), complete cleft palate (HP:0000175), hypertelorism (HP:0000316) and shawl scrotum (HP:0000049) (Table 1). Three heterozygous novel variants including *NAGLU*:c.311C>A (rank 1), *TGFB3*:c.785G>T (rank 2) and *KMT2D*:c.1372A>C (rank 3) were shortlisted by the filtering strategy (Table 3).

Heterozygous mutations of *NAGLU* have been identified to underlie Charcot-Marie-Tooth disease type 2V (CMT2V, OMIM # 616491). Patients with pathogenic *NAGLU* mutations present with a range of neurologic conditions including peripheral neuropathy and sensory impairment. The disease has a late age of onset and usually occurs between 18 and 61 years of age [49]. Considering the lack of phenotypic similarity between the patient *CL027* main features and CMT2V, it is unlikely that *NAGLU*:c.311C>A (Ψ_*i*_=8) underlies the patient’s condition.

Heterozygous mutations of *TGFB3* have been linked to two autosomal dominant conditions known as Arrhythmogenic right ventricular dysplasia 1 (ARVD1, OMIM # 107970) and Loeys-Dietz syndrome 5 (LDS5, # 615582). Clinical presentation in ARVD1 restricts to cardiovascular complications [50] whereas in LDS5 patients present cleft palate in combination with a range of cardiovascular complications including aortic aneurysms [51]. Considering that the proband *CL027* does not appear to have cardiovascular complications and given the lack of phenotypic similarity to the syndromes described above, it is highly unlikely that the *TGFB3*:c.785G>T (Ψ_*i*_=5.5) is causal in the context of the patient’s condition. Taken together, we could not conclusively resolve the molecular diagnosis in this patient.

#### Patient CL0331

The proband is a 10-year-old male patient from a non-consanguineous marriage. The patient present with normal development, unilateral cleft lip (HP:0100333), complete cleft palate (HP:0000175), hypertelorism (HP:0000316), telecanthus (HP:0000506), shawl scrotum (HP:0000049) and bilateral clinodactyly of the 5th finger (HP:0004209) (Table 1). Two variants including the NIPBL:c.7801A>G (rank 2) and *KIRREL3*:c.544A>T (rank 3) revealed to have high quality IGV reads.

Heterozygous mutations of *NIPBL* are identified to underlie autosomal dominant Cornelia de Lange syndrome-1 (CDLS1, OMIM # 122470) [52]. A wide clinical variability for CDLS1 has been described. The CDLS1 patients typically present with multisystem malformations including facial dysmorphisms, upper limb anomalies, developmental delay and mental retardation [53]. Twenty per cent of children with the CDLS1 present with cleft palate [54] and upper limb anomalies and the fifth finger clinodactyly is reported in 74% of cases [55]. Genitourinary anomalies including hypoplastic genitalia and cryptorchidism are fairly common among CDLS1 patients and are reported in 57% and 73% of the male patients respectively [55,56]. Lalatta *et aI*. [57] reported hypertelorism in a neonatal case of CDLS1 with novel mutation at exon
35 of *NIPBL*. The wide variability in the clinical presentations of the disease renders its diagnosis challenging. Severe forms of CDLS1 are readily diagnosed at an early age (>90% below the age of 2), but diagnosis becomes more challenging as patients grow older [53]. Establishing a diagnosis in the milder form of CDLS1 is even more challenging as patients do not express the characteristic facial appearance of the disease until 2 to 3 years of age. The facial features of CDLS1 in patients with milder form gradually fade as patients grow older and it becomes less significant after age nine years [58]. Given the variability in the clinical expression of the disease and extensive phenotypic similarity between the proband’s main features and CDLS1, it is highly likely that non-synonymous NIPBL:c.7801A>G on exon 45 underlies the patient’s condition.

#### Patient CL035

The patient is an 8-month-old infant of a non-consanguineous marriage. The proband’s main features include developmental delay (HP: 0011342), unilateral cleft lip (HP: 0100333) and complete cleft palate (HP: 0000175) 3). The phenotypic features reported for this patient are very broad and unspecific and therefore establishing a molecular diagnosis is challenging. Among the variants prioritised in this patient, only the GRIN2A:c.G136A fulfilled the quality criteria. Mutations in *GRIN2A* have been identified to cause variable neurodevelopmental phenotypes including the autosomal dominant focal epilepsy with speech disorder (FESD, #p 245570). Endele *et al*. [59] reported two patients with mutation in *GRIN2A* who present with facial dysmorphisms and neurodevelopmental delay. The facial characteristic of one patient with translocation breakpoint disrupting GRIN2A restrict to the short nose, but the nature of facial dysmorphism in another patient with non-synonymous GRIN2A:c.652C>T mutation is not fully described [59]. Considering incomplete penetrance and variable phenotype reported in the context of *GRIN2A* mutations [60,61], it is possible that identified non-synonymous mutation at *GRIN2A:* c.136G>A (Ψ_*i*_= 4) is causal in the patient. The FESD patients usually present with childhood-onset seizures that may last until adulthood [62]. Considering that phenotyping in the proband *CL035* was carried out during infancy further neuropsychological assessment is required in order to establish a diagnosis.

#### Patient CL036

The patient is a five years old boy of a non-consanguineous marriage present with developmental delay (HP:0011342), unilateral cleft lip (HP:0100333), complete cleft palate (HP:0000175), hypertelorism (HP:0000316), telecanthus (HP:0000506), shawl scrotum (HP:0000049), cryptorchidism (HP:0000028), clinodactyly of the 5th finger (HP:0030084) and heart murmur (HP:0030148) (Table 1). The patient had a low height, and weight percentile for his age and proband’s maternal cousin was also present with CLP. Five novel variants were identified as putatively pathogenic (Table 3). Investigation of genotype quality scores in parallel with visual inspection of read alignments for shortlisted variants revealed that *SMARCAĄ*: c.889A>C (rank 4 Ψ_*i*_= 6, DP = 27, GQ = 31) and *ARID1B*:c.293A>C (rank 5, Ψ_*i*_= 3, DP = 22, GQ = 28) are probably spurious calls and therefore they were excluded from further analysis. The genotype quality score for the non-synonymous *PTPN11*: c.323C>A (rank 1) variant was also very low (GQ = 28) and considering the low depth of coverage at this site (DP = 11) a reliable basis for a heterozygous call could not be established.

Two variants including *KMT2D*: c.16493C>T (rank 2, ψ = 7) and *BRCA2*: c.9425A>T (rank 3, Ψ_*i*_ = 7) were identified to have high read depth and high genotype quality (DP > 80 and GQ = 99) for confident variant analysis.

The *KMT2D* gene encodes a histone methyltransferase that is an important component of ASCOM complex which has been shown to regulate the transcriptional activity of beta-globin and estrogen receptor genes [63,64]. Heterozygous mutations of *KMT2D* are identified to underlie autosomal dominant Kabuki syndrome 1 (KABUK1, OMIM # 147920) [65]. Individuals with KABUK1 syndrome present with multiple congenital anomalies and characteristic facial features that typically include highly arched and sparse eyebrows, depressed nasal tip, short columella and large earlobes [66]. Cleft lip and/or cleft palate have been reported in 35% of cases [34]. Genitourinary abnormalities including cryptorchidism and micropenis have been observed in approximately 25% and 10% of male patients, respectively [34, 67]. Clinodactyly of the fifth digit is one of the major skeletal anomalies reported in 50% of patients [67]. Additional phenotypes of the patient that are consistent with KABUK1 syndrome include developmental delay and, heart murmur. Developmental delay and variable degrees of mental retardation have been described in the context of KABUK1 syndrome [68]. Mild and moderate developmental delay collectively present in 67% of Kabuki patients [69] and cardiovascular anomalies have been reported in 42% of cases [67]. The cause of heart murmur in the proband *CL036* is not defined, but common heart defects (including Atrial septal defect (ASD), Ventricular septal defect (VSD) and Patent ductus arteriosus (PDA)) that result in childhood heart murmur have been identified in 90.6% of Kabuki patients diagnosed prenatally or at an early age [70]. Given the extensive phenotypic similarity between the patient’s main features and KABUK1 syndrome and, also high combined rank score for the *KMT2D*: c.16493C>T mutation (rank 2, Ψ_*i*_ = 7), it is possible that this variant underlies the patient condition. Evaluation of the phenotypic features, revealed from patients photos, by three independent clinical experts at the Southampton University Hospital strongly supported the diagnosis of Kabuki syndrome.

The *BRCA2* gene was primarily included in our tiered gene list since transcriptional dysregulation of *BRCA2* are identified to associate with susceptibility to DNA damage and non-syndromic CLP [71]. Given the absence of direct link between *BRCA2* mutations and the patient’s main features, it is possible that the *BRCA2*: c.9425A>T is an incidental finding without any implication on the patient’s phenotype. Although this variant appears unlikely to have any impact on patient’s health, because of the *BRCA2* role in familial Breast and ovarian cancer (OMIM #600185), the identified mutation might have a clinical significance for female relatives of the patient. Therefore in compliance with the ACMG guideline for reporting secondary findings [72], it has been brought to the attention of recruiting clinician as an incidental finding.

#### Patient CL039

The patient is an 8-year-old boy from a non-consanguineous parent. The proband’s sister presented with microtia, and there is no history of CLP in his family. The patient’s main features include hyperactivity disorder (HP:0000752), unilateral cleft lip (HP:0100333), complete cleft palate (HP:0000175), hypertelorism (HP:0000316) and shawl scrotum (HP:0000049) (Table 1). Variant analysis shortlisted six novel mutations in genes including *CREBBP, OFD1, TBXį, ATR, ANKRD11* and *SALL4* (Table 3). Inspection of read alignments in IGV in parallel with the investigation of genotype quality scores revealed that only the ATR:c.3025G>A (rank 4, Ψ_*i*_ = 2.5, DP = 56, GQ= 99) appears to be a reliable call. The remaining variants reflect alignment errors and therefore were excluded from further analysis.

Heterozygous mutations of the *ATR* gene have been identified to underlie autosomal dominant cutaneous telangiectasia and cancer syndrome (FCTCS, OMIM # 614564) [73]. Patients with pathogenic heterozygous mutations of *ATR* present with cutaneous telangiectasia and dispersed alopecia during infancy [74]. Given the lack of reported telangiectasies complications in the proband *CL039*, it is unlikely that heterozygous ATR:c.3025G>A is related to the patient’s condition.

Homozygous or compound heterozygous mutations of ATR have been reported to cause autosomal recessive Seckel syndrome 1 (SCKL1, OMIM # 210600). Among phenotypic features reported for the proband *CL039*, cleft lip/cleft palate and hyperactivity have also been documented in SCKL1 patients [75-77]. Considering that no other non-synonymous mutation was identified in the ATR gene, the patient’s condition could not be linked to Seckel syndrome 1. Taken together variant analysis in the patient *CL039* failed to identify the causal variant and molecular diagnosis in this patient remained unresolved.

## Discussion

Despite the fact that molecularly confirmed cases of AAS have been entirely attributed to the pathogenic mutations of *FGD1*, we did not identify any variant with the pathogenic implication in this gene across the samples analysed. Failure to identify *FGD1* mutations could be attributed to several factors that are discussed below.

Aarskog-Scott syndrome is a rare disorder with extensive phenotypic similarity with other rare phenotypes [8]. Genotype-phenotype correlation of the disease is not fully understood, and molecular diagnosis in only 20% of cases is resolved [8]. Differential diagnosis when molecular aetiology of the disease is inconclusive proved to be challenging. Given the overlapping nature of facial characteristics in rare disorders [78], the possibility of alternative diagnoses compatible with molecular findings must be considered. Because of this, it is possible that variants identified in the five patients might genuinely be related to the patients’ phenotype and underlie their condition.

Secondly, the TruSight One capture kit does not provide a uniform capture across the 18 exons of the *FGD1* gene (Figure 1). To investigate whether the failure to identify *FGD1* mutation in this study is due to sample quality or inherent deficiency of TruSight One kit for capturing *FGD1* coding region, normalised read counts across the gene for the eight samples that passed alignment QC compared to 18 controls from the same sequencing batch. Although coverage efficiency significantly differs between exons of the *FGD1* (p< 0.001, One-way ANNOVA), capture coverage of the gene for AAS samples is not significantly different between the AAS cases and controls (*p* = 0.9998, One-tailed t-test) (Figure 2). Therefore a possible explanation for the failure to identify *FGD1* mutations can be the limitations of TruSight One kit for capturing *FGD1* exons. In particular exons, 5, 10 and 16 were identified to have low capture coverage in the TruSight One kit.

Investigation of mutational spectrum in the coding region of *FGD1* revealed that the highest number of novel SNVs map to the exon six but the highest density of mutational events *(i.e*. number of SNVs per length of the exon) occurs at the exon 13 (Figure 3). The depth of coverage at both exons was sufficient for variant calling in our study. Only one variant have been identified on each of the exon 5, and exon 10 and no variant have been reported in the exon 16 (Figure 3). Total number of SNVs identified in these three exons constitute a mere 5.74% of the total mutational density across the coding length of *FGD1* gene, and therefore even with a reduced capture in these exons, we have still been able to robustly screen for pathogenic mutations in the remaining exons with theoretical 94.25% mutational density. However, given the modest statistical power of targeted exome sequencing for identifying *de-novo* heterozygous variants [79], the implication of low capture coverage in these exons for detecting *FGD1* variants merits further investigation.

Thirdly, the inherent deficiency of targeted exome sequencing in capturing sequences outside the coding region might also explain the lack of *FGD1* positive cases in our study. The *FGD1* intronic variants (including splicing SNVs) have been reported to underlie AAS [11,80]. Considering that targeted exome sequencing is ill-suited for identifying variants outside the coding region, we might have missed pathogenic variants in the intronic region.

Finally, the mean insert size for the nine samples analysed was ~143 bp. Low quality DNA or reduced mass input of gDNA changes the distribution of library fragments and result in smaller insert sizes than expected [81]. These smaller fragments are prone to exclusion during the subsequent clean-up step and result in a skewed fragment distribution. In view of this, skewed insert length clearly explains the unevenness in coverage distribution in our samples, especially for the *FGD1* gene. In addition, larger read lengths (≥ 170) are identified to provide a more uniform coverage distribution across the genes targeted and reduce the false negative rate in the variant detection by two fold [82]. Although larger read length is achieved at the expense of reduced depth, evenness in the coverage dramatically increases the diagnostic power of exome sequencing [82]. The average size of exons in the human genome is 170 bp, and 80-85% of exons are identified to be <200 bp in size [83,84]. Consequently, read lengths above 170bp provide an ideal size distribution for capturing not only exonic variants but also intronic variants that have implications in splicing. Because of this, skewed read length size in our study might provide an explanation for the reduced diagnostic power to identify the *FGD1* mutations in our samples. Given uneven coverage across the exons of *FGD1* in the TruSigth one panel and the reduced insert sizes in this study, screening of the *FGD1* gene in higher uniform coverage must be prioritised for unresolved cases.

While rare disorders are believed to be primarily caused by a single penetrant protein truncating mutation, there are cases in which patients with the same genetic defect present different phenotypes [85]. This variable expressivity is generally attributed to the role of modifier genes that collectively impact the penetrance, dominance, and expressivity of the mutation [86]. Besides, many rare disorders share extensive phenotypic similarity. The lack of granularity in phenotypic manifestations of rare disorders is particularly problematic when it comes to molecular diagnosis. Also there are occasions in which mutations in different genes can result in similar phenotypes [87]. The issue pertaining to the lack of consistency in phenotypic manifestations is more pronounced in the context of neurodevelopmental disorders [88], mainly because of a large number of genes identified to implicate in the disease pathogenicity [89]. In the absence of detailed phenotypic information, similarities between patients cannot be statistically tested, and the power of NGS analysis is reduced. Obtaining complete and update phenotypic information from patients residing in remote areas in Colombia presented a significant challenge in this study. Indeed, mixed phenotypic information with a variable degree of details and completeness appeared to be the main limitation for molecular diagnosis in these patients. Despite the provisional diagnosis of AAS, many features of the patients are unspecific to AAS. Clinically, this implicates that patients studied here perhaps present a heterogeneous group of disorders that share some phenotypic similarities including CLP, shawl scrotum and hypertelorism (the three most frequent features in the cohort). Interestingly, identification of plausible mutations in five individuals (out of the eight patients satisfied QC criteria) presents a strong case for alternative diagnosis across these patients.

To summarise, we applied targeted exome sequencing to 14 patients with the provisional diagnosis of Aarskog-Scott syndrome. Six samples were discarded from variant analysis early on as they did not comply with the quality standards required for confident variant analysis. Investigation of novel variants in the remaining eight samples resulted in the identification of putative causal variants for five samples including CL021, CL025, CL033, CL035 and CL036 (Table 4).

**Table 4.** Summary of putatively causa1 variants identified in ñve samples (Ψ_*i*_: Combined rank score; GQ: Genotype quality score)

| Sample | Variant | Exon | $\Psi_i$ | Depth | GQ |
| --- | --- | --- | --- | --- | --- |
| CL021 | SRCAP:c.A8036T | 34 | 5 | 166 | 99 |
| CL025 | OFD1:c.G2610C | 20 | 4.5 | 42 | 36 |
| CL033 | NIPBL:c.A7801G | 45 | 3.5 | 148 | 99 |
| CL035 | GRIN2A:c.G136A | 2 | 4 | 252 | 99 |
| CL036 | KMT2D:c.C16493T | 53 | 7 | 119 | 99 |

An important finding from this analysis is the Kabuki syndrome (KS) diagnosis in the patient *CL036*. Having reviewed the findings with our local clinical experts, we confirmed the Kabuki diagnosis in this patient. While KS was believed to have a higher incidence in Japanese populations, recent findings in other ethnic groups have extended the phenotypic spectrum of the disease and revealed new pathological mutations [90,91].

Establishing molecular diagnosis in the remaining patients requires further follow-up including in-depth investigation of phenotypic details that may corroborate the molecular finding from WES analysis.

## Ethics approval

Analysis of patients detailed in this study has been approved by the Research Ethics Committee at the Universidad de La Sabana, Bogota, Colombia (Comité de Ética en Investigación-Acta number: 29, 25 May 2012).

## References

1. Andrew C. Lidral, Lina M. Moreno, and Steven A. Bullard. Genetic Factors and Orofacial Clefting. Seminars in Orthodontics, 2008.

2. Liliana Arias Uruena, Ignacio Briceno Balcazar, Julio Martinez Lozano, Andrew Collins, and Daniel Alfredo Uricoechea Patino. Clinical Aspects associated with Syndromic forms of Orofacial Clefts in a Colombian population. Colombia medica (Cali, Colombia), 2015.

3. Astanand Jugessur, Fedik Rahimov, Rolv T. Lie, Allen J. Wilcox, Håkon K. Gjessing, Roy M. Nilsen, Truc Trung Nguyen, and Jeffrey C. Murray. Genetic variants in IRF6 and the risk of facial clefts: Single-marker and haplotype-based analyses in a population-based case-control study of facial clefts in Norway. Genetic Epidemiology, 2008.

4. Michael J. Dixon, Mary L. Marazita, Terri H. Beaty, and Jeffrey C. Murray. Cleft lip and palate: Understanding genetic and environmental influences, 2011.

5. Theresa M Zucchero, Margaret E Cooper, Brion S Maher, Sandra Daack-Hirsch, Buena Nepomuceno, Lucilene Ribeiro, Diana Caprau, Kaare Christensen, Yasushi Suzuki, Junichiro Machida, Nagato Natsume, Koh-Ichiro Yoshiura, Alexandre R Vieira, Ieda M Orioli, Eduardo E Castilla, Lina Moreno, Mauricio Arcos-Burgos, Andrew C Lidral, L Leigh Field, You-e Liu, Ajit Ray, Toby H Goldstein, Rebecca E Schultz, Min Shi, Marla K Johnson, Shinji Kondo, Brian C Schutte, Mary L Marazita, and Jeffrey C Murray. Interferon regulatory factor 6 (IRF6) gene variants and the risk of isolated cleft lip or palate. The New England journal of medicine, 2004.

6. A S Teebi, K K Naguib, S Al-Awadi, and Q A Al-Saleh. New autosomal recessive faciodigitogenital syndrome. Journal of Medical Genetics, 25(6):400–406, 6 1988.

7. Mariana Pérez-Coria, José J Lugo-Trampe, Michell Zamudio-Osuna, Iram P Rodríguez-Sánchez, Angel Lugo-Trampe, Beatriz de la Fuente-Cortez, Luis D Campos-Acevedo, and Laura E Martínez-de Villarreal. Identification of novel mutations in Mexican patients with Aarskog–Scott syndrome. Molecular Genetics and Genomic Medicine, 3(3):197–202, 2015.

8. M. Reza Jabalameli, Ignacio Briceno, Julio Martinez, Ignacio Briceno, Reuben J. Pengelly, Sarah Ennis, and Andrew Collins. Aarskog-Scott syndrome: phenotypic and genetic heterogeneity. AIMS Genetics, 3(1):49–59, 2016.

9. L Logie and M Porteous. Intelligence and development in Aarskog syndrome. Archives of Disease in Childhood, 79(4):359–360, 10 1998.

10. Alfredo Orrico, Lucia Galli, Jill Clayton-Smith, and Jean-Pierre Fryns. Clinical utility gene card for: Aarskog–Scott syndrome (faciogenital dysplasia). European Journal of Human Genetics, 19(11), 11 2011.

11. A. Orrico, L. Galli, L. Faivre, J. Clayton-Smith, S. M. Azzarello-Burri, J. M. Hertz, S. Jacquemont, R. Taurisano, I. Arroyo Carrera, E. Tarantino, K. Devriendt, D. Melis, T. Thelle, U. Meinhardt, and V. Sorrentino. Aarskog-Scott syndrome: Clinical update and report of nine novel mutations of the FGD1 gene. American Journal of Medical Genetics, Part A, 152(2):313–318, 2010.

12. Peter D. Stenson, Matthew Mort, Edward V. Ball, Katy Shaw, Andrew D. Phillips, and David N. Cooper. The Human Gene Mutation Database: Building a comprehensive mutation repository for clinical and molecular genetics, diagnostic testing and personalized genomic medicine, 2014.

13. Joanna S. Amberger, Carol A. Bocchini, Françcois Schiettecatte, Alan F. Scott, and Ada Hamosh. OMIM.org: Online Mendelian Inheritance in Man (OMIM), an Online catalog of human genes and genetic disorders. Nucleic Acids Research, 43(D1):D789–D798, 2015.

14. Heng Li, Bob Handsaker, Alec Wysoker, Tim Fennell, Jue Ruan, Nils Homer, Gabor Marth, Goncalo Abecasis, and Richard Durbin. The Sequence Alignment/Map format and SAMtools. Bioinformatics, 25(16):2078–2079, 2009.

15. Geraldine A. Van der Auwera, Mauricio O. Carneiro, Christopher Hartl, Ryan Poplin, Guillermo del Angel, Ami Levy-Moonshine, Tadeusz Jordan, Khalid Shakir, David Roazen, Joel Thibault, Eric Banks, Kiran V. Garimella, David Altshuler, Stacey Gabriel, and Mark A. DePristo. From fastQ data to high-confidence variant calls: The genome analysis toolkit best practices pipeline. Current Protocols in Bioinformatics, (SUPL.43), 2013.

16. Kai Wang, Mingyao Li, and Hakon Hakonarson. ANNOVAR: Functional annotation of genetic variants from high-throughput sequencing data. Nucleic Acids Research, 38(16), 2010.

17. S. T. Sherry. dbSNP: the NCBI database of genetic variation. Nucleic Acids Research, 29(1):308–311, 2001.

18. The 1000 Genomes Project Consortium. A global reference for human genetic variation, 2015.

19. Monkol Lek, Konrad J Karczewski, Eric V Minikel, Kaitlin E Samocha, Eric Banks, Timothy Fennell, Anne H O’Donnell-Luria, James S Ware, Andrew J Hill, Beryl B Cummings, Taru Tukiainen, Daniel P Birnbaum, Jack A Kosmicki, Laramie E Duncan, Karol Estrada, Fengmei Zhao, James Zou, Emma Pierce-Hoffman, Joanne Berghout, David N Cooper, Nicole Deflaux, Mark DePristo, Ron Do, Jason Flannick, Menachem Fromer, Laura Gauthier, Jackie Goldstein, Namrata Gupta, Daniel Howrigan, Adam Kiezun, Mitja I Kurki, Ami Levy Moonshine, Pradeep Natarajan, Lorena Orozco, Gina M Peloso, Ryan Poplin, Manuel A Rivas, Valentin Ruano-Rubio, Samuel A Rose, Douglas M Ruderfer, Khalid Shakir, Peter D Stenson, Christine Stevens, Brett P Thomas, Grace Tiao, Maria T Tusie-Luna, Ben Weisburd, Hong-Hee Won, Dongmei Yu, David M Altshuler, Diego Ardissino, Michael Boehnke, John Danesh, Stacey Donnelly, Roberto Elosua, Jose C Florez, Stacey B Gabriel, Gad Getz, Stephen J Glatt, Christina M Hultman, Sekar Kathiresan, Markku Laakso, Steven McCarroll, Mark I McCarthy, Dermot McGovern, Ruth McPherson, Benjamin M Neale, Aarno Palotie, Shaun M Purcell, Danish Saleheen, Jeremiah M Scharf, Pamela Sklar, Patrick F Sullivan, Jaakko Tuomilehto, Ming T Tsuang, Hugh C Watkins, James G Wilson, Mark J Daly, Daniel G MacArthur, and Exome Aggregation Consortium. Analysis of protein-coding genetic variation in 60,706 humans. Nature, 536(7616):285–291, 8 2016.

20. Karthik A Jagadeesh, Aaron M Wenger, Mark J Berger, Harendra Guturu, Peter D Stenson, David N Cooper, Jonathan A Bernstein, and Gill Bejerano. M-CAP eliminates a majority of variants of uncertain significance in clinical exomes at high sensitivity. Nature genetics, 48(12):1581–1586, 2016.

21. Ivan Adzhubei, Daniel M. Jordan, and Shamil R. Sunyaev. Predicting functional effect of human missense mutations using PolyPhen-2. Current Protocols in Human Genetics, (SUPPL.76), 2013.

22. Pauline C. Ng and Steven Henikoff. Predicting the Effects of Amino Acid Substitutions on Protein Function. Annual Review of Genomics and Human Genetics, 7(1):61–80, 2006.

23. Sung Chun and Justin C. Fay. Identification of deleterious mutations within three human genomes. Genome Research, 19(9):1553–1561, 2009.

24. Hashem A. Shihab, Julian Gough, David N. Cooper, Peter D. Stenson, Gary L A Barker, Keith J. Edwards, Ian N M Day, and Tom R. Gaunt. Predicting the Functional, Molecular, and Phenotypic Consequences of Amino Acid Substitutions using Hidden Markov Models. Human Mutation, 34(1):57–65, 2013.

25. Chengliang Dong, Peng Wei, Xueqiu Jian, Richard Gibbs, Eric Boerwinkle, Kai Wang, and Xiaoming Liu. Comparison and integration of deleteriousness prediction methods for nonsynonymous SNVs in whole exome sequencing studies. Human Molecular Genetics, 24(8):2125–2137, 2015.

26. Martin Kircher, Daniela M Witten, Preti Jain, Brian J O’Roak, Gregory M Cooper, Jay Shendure, Brian J O Roak, Gregory M Cooper, and Jay Shendure. A general framework for estimating the relative pathogenicity of human genetic variants. Nature Genetics, 46(3):310–315, 2014.

27. Gregory M. Cooper, Eric A. Stone, George Asimenos, Eric D. Green, Serafim Batzoglou, and Arend Sidow. Distribution and intensity of constraint in mammalian genomic sequence. Genome Research, 2005.

28. Eugene V. Davydov, David L. Goode, Marina Sirota, Gregory M. Cooper, Arend Sidow, and Serafim Batzoglou. Identifying a high fraction of the human genome to be under selective constraint using GERP++. PLoS Computational Biology, 6(12), 2010.

29. William McLaren, Laurent Gil, Sarah E. Hunt, Harpreet Singh Riat, Graham R. S. Ritchie, Anja Thormann, Paul Flicek, and Fiona Cunningham. The Ensembl Variant Effect Predictor. Genome Biology, 17(1):122, 2016.

30. François-Olivier Desmet, Dalil Hamroun, Marine Lalande, Gwenaëlle Collod-Béroud, Mireille Claustres, and Christophe Béroud. Human Splicing Finder: an online bioinformatics tool to predict splicing signals. Nucleic Acids Research, 37(9):e67–e67, 5 2009.

31. Rebecca L. Hood, Matthew A. Lines, Sarah M. Nikkel, Jeremy Schwartzentruber, Chandree Beaulieu, Małgorzata J.M. Nowaczyk, Judith Allanson, Chong Ae Kim, Dagmar Wieczorek, Jukka S. Moilanen, Didier Lacombe, Gabriele Gillessen-Kaesbach, Margo L. Whiteford, Caio Robledo D.C. Quaio, Israel Gomy, Debora R. Bertola, Beate Albrecht, Konrad Platzer, George McGillivray, Ruobing Zou, D. Ross McLeod, Albert E. Chudley, Bernard N. Chodirker, Janet Marcadier, Jacek Majewski, Dennis E. Bulman, Susan M. White, and Kym M. Boycott. Mutations in SRCAP, encoding SNF2-related CREBBP activator protein, cause Floating-Harbor syndrome. American Journal of Human Genetics, 90(2):308–313, 2012.

32. M A Monroy, D D Ruhl, X Xu, D K Granner, P Yaciuk, and J C Chrivia. Regulation of cAMP-responsive element-binding protein-mediated transcription by the SNF2/SWI-related protein, SRCAP. The Journal of biological chemistry, 276(44):40721–6, 11 2001.

33. Keisuke Nagasaki, Tadashi Asami, Hidetoshi Sato, Yohei Ogawa, Toru Kikuchi, Akihiko Saitoh, Tsutomu Ogata, and Maki Fukami. Long-term follow-up study for a patient with Floating-Harbor syndrome due to a hotspot SRCAP mutation. American Journal of Medical Genetics, Part A, 164(3):731–735, 2014.

34. Margaret P. Adam and L. Hudgins. Kabuki syndrome: A review, 2005.

35. Dominique Well, Stéphane Blanchard, Josseline Kaplan, Parry Guilford, Fernando Gibson, James Walsh, Philomena Mburu, Anabel Varela, Jacqueline Levilliers, Michael Weston, D., Phillip Kelley, M., William Kimberling, J., Mariette Wagenaar, Fabienne Levi-Acobas, Dominique Larget-Piet, Arnold Munnich, Karen Steel, P., Steve D. Brown, M., and Christine Petit. Defective myosin VIIA gene responsible for Usher syndrome type IB. Nature, 374(6517):60–61, 1995.

36. Yi Sun, Jing Chen, Hanjun Sun, Jing Cheng, Jianzhong Li, Yu Lu, Yanping Lu, Zhanguo Jin, Yuhua Zhu, Xiaomei Ouyang, Denise Yan, Pu Dai, Dongyi Han, Weiyan Yang, Rongguang Wang, Xuezhong Liu, and Huijun Yuan. Novel missense mutations in MYO7A underlying postlingual high-or low-frequency non-syndromic hearing impairment in two large families from China. Journal of human genetics, 56(1):64–70, 2011.

37. Jessica W Witherspoon and Katherine G Meilleur. Review of RyR1 pathway and associated pathomechanisms. Acta neuropathologica communications, 4(1):121, 2016.

38. Diana Xerxes Bharucha-Goebel, Mariarita Santi, Livija Medne, Kristin Zukosky, Jahannaz Dastgir, Perry B. Shieh, Thomas Winder, Gihan Tennekoon, Richard S. Finkel, James J. Dowling, Nicole Monnier, and Carsten G. Bonnemann. Severe congenital RYRI-associated myopathy: The expanding clinicopathologic and genetic spectrum. Neurology, 80(17):1584–1589, 2013.

39. Hyam Isaacs and Margaret E. Badenhorst. Dominantly inherited malignant hyperthermia (MH) in the King-Denborough Syndrome. Muscle and Nerve, 15(6):740–742, 1992.

40. Rachel Robinson, Danielle Carpenter, Marie Anne Shaw, Jane Halsall, and Philip Hopkins. Mutations in RYR1 in malignant hypertheraiia and central core disease, 2006.

41. Lewis C Cantley. The phosphoinositide 3-kinase pathway. Science (New York, N.Y.), 296(5573):1655–7, 2002.

42. Clea Bárcena, Víctor Quesada, Annachiara De Sandre-Giovannoli, Diana a Puente, Joaquin Fernández-Toral, Sabine Sigaudy, Anwar Baban, Nicolas Lévy, Gloria Velasco, and Carlos López-Otín. Exome sequencing identifies a novel mutation in PIK3R1 as the cause of SHORT syndrome. BMC medical genetics, 15(1):51, 2014.

43. David A. Dyment, Amanda C. Smith, Diana Alcantara, Jeremy A. Schwartzentruber, Lina Basel-Vanagaite, Cynthia J. Curry, I. Karen Temple, William Reardon, Sahar Mansour, Mushfequr R. Haq, Rodney Gilbert, Ordan J. Lehmann, Megan R. Vanstone, Chandree L. Beaulieu, Jacek Majewski, Dennis E. Bulman, Mark O’Driscoll, Kym M. Boycott, and A. Micheil Innes. Mutations in PIK3R1 cause SHORT syndrome. American Journal of Human Genetics, 93(1):158–166, 2013.

44. Raoul C M Hennekam. Rubinstein-Taybi syndrome. European journal of human genetics: EJHG, 14(9):981–985, 2006.

45. R C Hennekam and J M Van Doorne. Oral aspects of Rubinstein-Taybi syndrome. American journal of medical genetics. Supplement, 6:42–47, 1990.

46. Margarita Larralde de Luna, María Luisa Raspa, and Jesús Ibargoyen. Oral-Facial-Digital Type 1 Syndrome of Papillon-Lóeage and Psaume. Pediatric Dermatology, 9(1):52–56, 3 1992.

47. M I Ferrante, G Giorgio, S A Feather, A Bulfone, V Wright, M Ghiani, A Selicorni, L Gammaro, F Scolari, A S Woolf, O Sylvie, L Bernard, S Malcolm, R Winter, A Ballabio, and B Franco. Identification of the gene for oral-facial-digital type I syndrome. Am J Hum Genet, 68(3):569–576, 2001.

48. Y. Tsurusaki, T. Kosho, K. Hatasaki, Y. Narumi, K. Wakui, Y. Fukushima, H. Doi, H. Saitsu, N. Miyake, and N. Matsumoto. Exome sequencing in a family with an X-linked lethal malformation syndrome: Clinical consequences of hemizygous truncating OFD1 mutations in male patients. Clinical Genetics, 83(2):135–144, 2013.

49. Martine Tétreault, Michael Gonzalez, Marie Josée Dicaire, Pierre Allard, Kalle Gehring, Diane Leblanc, Nadine Leclerc, Ronald Schondorf, Jean Mathieu, Stephan Zuchner, and Bernard Brais. Adult-onset painful axonal polyneuropathy caused by a dominant NAGLU mutation. Brain, 138(6):1477–1483, 2015.

50. Cristina Basso, Domenico Corrado, Barbara Bauce, and Gaetano Thiene. Arrhythmogenic right ventricular cardiomyopathy. Circulation: Arrhythmia and Electrophysiology, 5(6):1233–1246, 2012.

51. Aida M. Bertoli-Avella, Elisabeth Gillis, Hiroko Morisaki, Judith M A Verhagen, Bianca M. de Graaf, Gerarda van de Beek, Elena Gallo, Boudewijn P T Kruithof, Hanka Venselaar, Loretha A. Myers, Steven Laga, Alexander J. Doyle, Gretchen Oswald, Gert W A van Cappellen, Itaru Yamanaka, Robert M. van der Helm, Berna Beverloo, Annelies de Klein, Luba Pardo, Martin Lammens, Christina Evers, Koenraad Devriendt, Michiel Dumoulein, Janneke Timmermans, Hennie T. Bruggenwirth, Frans Verheijen, Inez Rodrigus, Gareth Baynam, Marlies Kempers, Johan Saenen, Emeline M. Van Craenenbroeck, Kenji Minatoya, Ritsu Matsukawa, Takuro Tsukube, Noriaki Kubo, Robert Hofstra, Marie J ose Goumans, Jos A. Bekkers, Jolien W. Roos-Hesselink, Ingrid M B H van de Laar, Harry C. Dietz, Lut Van Laer, Takayuki Morisaki, Marja W. Wessels, and Bart L. Loeys. Mutations in a TGF-β ligand, TGFB3, cause syndromic aortic aneurysms and dissections. Journal of the American College of Cardiology, 65(13):1324–1336, 2015.

52. Martine Isabel Boyle, Cathrine Jespersgaard, Karen Brøndum-Nielsen, Anne-Marie Bisgaard, and Zeynep Tümer. Cornelia de Lange Syndrome. Clinical genetics, (March):38–41, 2014.

53. Sarika Rohatgi, Dinah Clark, Antonie D Kline, Laird G Jackson, Juan Pie, Victoria Siu, Feliciano J Ramos, Ian D Krantz, and Matthew A Deardorff. Facial diagnosis of mild and variant CdLS: Insights from a dysmorphologist survey. American Journal of Medical Genetics Part A, 152A(7):1641–1653, 7 2010.

54. Antonie D. Kline, Marco Grados, Paul Sponseller, Howard P. Levy, Natalie Blagowidow, Christianne Schoedel, Joni Rampolla, Douglas K. Clemens, Ian Krantz, Amy Kimball, Carmen Pichard, and David Tuchman. Natural history of aging in Cornelia de Lange syndrome. American Journal of Medical Genetics, Part C: Seminars in Medical Genetics, 145(3):248–260, 2007.

55. L. Jackson, A. D. Kline, M. A. Barr, and S. Koch. de Lange syndrome: A clinical review of 310 individuals. In American Journal of Medical Genetics, volume 47, pages 940–946, 1993.

56. Antonie D. Kline, Ian D. Krantz, Annemarie Sommer, Mark Kliewer, Laird G. Jackson, David R. FitzPatrick, Alex V. Levin, and Angelo Selicorni. Cornelia de Lange syndrome: Clinical review, diagnostic and scoring systems, and anticipatory guidance, 2007.

57. Faustina Lalatta, Silvia Russo, Barbara Gentilin, Luigina Spaccini, Chiara Boschetto, Florinda Cavalleri, Maura Masciadri, Cristina Gervasini, Angela Bentivegna, Paola Castronovo, and Lidia Larizza. Prenatal/neonatal pathology in two cases of Cornelia de Lange syndrome harboring novel mutations of NIPBL. Genetics in Medicine, 9(3):188–194, 2007.

58. J E Allanson, R C Hennekam, and M Ireland. De Lange syndrome: subjective and objective comparison of the classical and mild phenotypes. Journal of medical genetics, 34(8):645–50, 1997.

59. S Endele, G Rosenberger, K Geider, B Popp, C Tamer, I Stefanova, M Milh, F Kortum, A Fritsch, F K Pientka, Y Hellenbroich, V M Kalscheuer, J Kohlhase, U Moog, G Rappold, A Rauch, H H Ropers, S von Spiczak, H Tonnies, N Villeneuve, L Villard, B Zabel, M Zenker, B Laube, A Reis, D Wieczorek, L Van Maldergem, and K Kutsche. Mutations in GRIN2A and GRIN2B encoding regulatory subunits of NMDA receptors cause variable neurodevelopmental phenotypes. Nat Genet, 42(11):1021–1026, 2010.

60. Gaetan Lesca, Gabrielle Rudolf, Nadine Bruneau, Natalia Lozovaya, Audrey Labalme, Nadia Boutry-Kryza, Manal Salmi, Timur Tsintsadze, Laura Addis, Jacques Motte, Sukhvir Wright, Vera Tsintsadze, Anne Michel, Diane Doummar, Karine Lascelles, Lisa Strug, Patrick Waters, Julitta de Bellescize, Pascal Vrielynck, Anne de Saint Martin, Dorothee Ville, Philippe Ryvlin, Alexis Arzimanoglou, Edouard Hirsch, Angela Vincent, Deb Pal, Nail Burnashev, Damien Sanlaville, and Pierre Szepetowski. GRIN2A mutations in acquired epileptic aphasia and related childhood focal epilepsies and encephalopathies with speech and language dysfunction. Nature genetics, 45(9):1061–6, 2013.

61. Johannes R Lemke, Dennis Lal, Eva M Reinthaler, Isabelle Steiner, Michael Nothnagel, Michael Alber, Kirsten Geider, Bodo Laube, Michael Schwake, Katrin Finsterwalder, Andre Franke, Markus Schilhabel, Johanna A Jähn, Hiltrud Muhle, Rainer Boor, Wim Van Paesschen, Roberto Caraballo, Natalio Fejerman, Sarah Weckhuysen, Peter De Jonghe, Jan Larsen, Rikke S Møller, Helle Hjalgrim, Laura Addis, Shan Tang, Elaine Hughes, Deb K Pal, Kadi Veri, Ulvi Vaher, Tiina Talvik, Petia Dimova, Rosa Guerrero López, Jose M Serratosa, Tarja Linnankivi, Anna-Elina Lehesjoki, Susanne Ruf, Markus Wolff, Sarah Buerki, Gabriele Wohlrab, Judith Kroell, Alexandre N Datta, Barbara Fiedler, Gerhard Kurlemann, Gerhard Kluger, Andreas Hahn, D Edda Haberlandt, Christina Kutzer, Jürgen Sperner, Felicitas Becker, Yvonne G Weber, Martha Feucht, Hannelore Steinböck, Birgit Neophythou, Gabriel M Ronen, Ursula Gruber-Sedlmayr, Julia Geldner, Robert J Harvey, Per Hoffmann, Stefan Herms, Janine Altmuller, Mohammad R Toliat, Holger Thiele, Peter Nürnberg, Christian Wilhelm, Ulrich Stephani, Ingo Helbig, Holger Lerche, Fritz Zimprich, Bernd A Neubauer, Saskia Biskup, and Sarah von Spiczak. Mutations in GRIN2A cause idiopathic focal epilepsy with rolandic spikes. Nature genetics, 45(9):1067–72, 2013.

62. Gemma L Carvill, Brigid M Regan, Simone C Yendle, Brian J O’Roak, Natalia Lozovaya, Nadine Bruneau, Nail Burnashev, Adiba Khan, Joseph Cook, Eileen Geraghty, Lynette G Sadleir, Samantha J Turner, Meng-Han Tsai, Richard Webster, Robert Ouvrier, John A Damiano, Samuel F Berkovic, Jay Shendure, Michael S Hildebrand, Pierre Szepetowski, Ingrid E Scheffer, and Heather C Mefford. GRIN2A mutations cause epilepsy-aphasia spectrum disorders. Nature genetics, 45(9):1073–1076, 2013.

63. Cyrus Martin and Yi Zhang. The diverse functions of histone lysine methylation. Nature Reviews Molecular Cell Biology, 6(11):838–49, 2005.

64. Eneda Toska, Hatice U. Osmanbeyoglu, Pau Castel, Carmen Chan, Ronald C. Hendrickson, Moshe Elkabets, Maura N. Dickler, Maurizio Scaltriti, Christina S. Leslie, Scott A. Armstrong, and Jose Baselga. PI3K pathway regulates ER-dependent transcription in breast cancer through the epigenetic regulator KMT2D. Science, 355(6331):1324–1330, 2017.

65. Sarah B Ng, Abigail W Bigham, Kati J Buckingham, Mark C Hannibal, Margaret J McMillin, Heidi I Gildersleeve, Anita E Beck, Holly K Tabor, Gregory M Cooper, Heather C Mefford, Choli Lee, Emily H Turner, Joshua D Smith, Mark J Rieder, Koh-Ichiro Yoshiura, Naomichi Matsumoto, Tohru Ohta, Norio Niikawa, Deborah A Nickerson, Michael J Bamshad, and Jay Shendure. Exome sequencing identifies MLL2 mutations as a cause of Kabuki syndrome. Nature genetics, 42(9):790–793, 2010.

66. Mark C. Hannibal, Kati J. Buckingham, Sarah B. Ng, Jeffrey E. Ming, Anita E. Beck, Margaret J. Mcmillin, Heidi I. Gildersleeve, Abigail W. Bigham, Holly K. Tabor, Heather C. Mefford, Joseph Cook, Koh Ichiro Yoshiura, Tadashi Matsumoto, Naomichi Matsumoto, Noriko Miyake, Hidefumi Tonoki, Kenji Naritomi, Tadashi Kaname, Toshiro Nagai, Hirofumi Ohashi, Kenji Kurosawa, Jia Woei Hou, Tohru Ohta, Deshung Liang, Akira Sudo, Colleen A. Morris, Siddharth Banka, Graeme C. Black, Jill Clayton-Smith, Deborah A. Nickerson, Elaine H. Zackai, Tamim H. Shaikh, Dian Donnai, Norio Niikawa, Jay Shendure, and Michael J. Bamshad. Spectrum of MLL2 (ALR) mutations in 110 cases of Kabuki syndrome. American Journal of Medical Genetics, Part A, 155(7):1511–1516, 2011.

67. Naomichi Matsumoto and Norio Niikawa. Kabuki make-up syndrome: a review. American journal of medical genetics. Part C, Seminars in medical genetics, 117C VN –(1):57–65, 2003.

68. H Kawame, M C Hannibal, L Hudgins, and R a Pagon. Phenotypic spectrum and management issues in Kabuki syndrome. The Journal of pediatrics, 134(4):480–5, 1999.

69. Maria Cristina Digilio, Bruno Marino, Alessandra Toscano, Aldo Giannotti, and Bruno Dallapiccola. Congenital heart defects in Kabuki syndrome. American Journal of Medical Genetics, 100(4):269–274, 5 2001.

70. Shi Min Yuan. Congenital heart defects in Kabuki syndrome, 2013.

71. Gerson Shigeru Kobayashi, Lucas Alvizi, Daniele Yumi Sunaga, Philippa Francis-West, Anna Kuta, Bruno Vinícius Pimenta Almada, Simone Gomes Ferreira, Leonardo Carmo de Andrade-Lima, Daniela Franco Bueno, Cáassio Eduardo Raposo-Amaral, Carlos Frederico Menck, and Maria Rita Passos-Bueno. Susceptibility to DNA Damage as a Molecular Mechanism for Non-Syndromic Cleft Lip and Palate. PLoS ONE, 8(6), 2013.

72. Sarah S Kalia, Kathy Adelman, Sherri J Bale, Wendy K Chung, Christine Eng, James P Evans, Gail E Herman, Sophia B Hufnagel, Teri E Klein, Bruce R Korf, Kent D McKelvey, Kelly E Ormond, C Sue Richards, Christopher N Vlangos, Michael Watson, Christa L Martin, and David T Miller. Recommendations for reporting of secondary findings in clinical exome and genome sequencing, 2016 update (ACMG SF v2.0): a policy statement of the American College of Medical Genetics and Genomics. Genetics In Medicine, 19:249, 11 2016.

73. Mark Ambrose and Richard A. Gatti. Pathogenesis of ataxia-telangiectasia: the next generation of ATM functions., 2013.

74. Akio Tanaka, Sarah Weinel, Nikoletta Nagy, Mark O’Driscoll, Joey E. Lai-Cheong, Carol L. Kulp-Shorten, Alfred Knable, Gillian Carpenter, Sheila A. Fisher, Makiko Hiragun, Yuhki Yanase, Michihiro Hide, Jeffrey Callen, and John A. McGrath. Germline mutation in ATR in autosomal– dominant oropharyngeal cancer syndrome, 2012.

75. Jyotsna Murthy, Krishna G. Seshadri, Padmasani Venkat Ramanan, Arvind Rajamani, and Altaf Hussain. A Case of Cleft Lip and Palate Associated with Seckel Syndrome. Cleft Palate-Craniofacial Journal, 41(2):202–205, 2004.

76. Arvind Rajamani, Vijaylakshmi Kamat, Jyotsna Murthy, and Syed Altaf Hussain. Anesthesia for cleft lip surgery in a child with Seckel syndrome - A case report. Paediatric Anaesthesia, 15(4):338–341, 2005.

77. Quarantelli M Martinelli P Paladini D Napolitano R Maruotti GM. Prenatal diagnosis of Seckel syndrome on 3-dimensional sonography and magnetic resonance imaging. Journal Of Ultrasound In Medicine (ISSN: 0278-Ą297, 1550-9613), 2(3):369–374, 2009.

78. Deciphering Developmental Disorders Study. Prevalence and architecture of de novo mutations in developmental disorders. Nature, 542(7642):433–438, 2017.

79. Degui Zhi and Rui Chen. Statistical guidance for experimental design and data analysis of mutation detection in rare monogenic Mendelian diseases by exome sequencing. PLoS ONE, 7(2), 2012.

80. Emmelien Aten, Yu Sun, Rowida Almomani, Gijs W E Santen, Tobias Messemaker, Saskia M. Maas, Martijn H. Breuning, and Johan T. Den Dunnen. Exome Sequencing Identifies A Branch Point Variant in Aarskog-Scott Syndrome. Human Mutation, 34(3):430–434, 2013.

81. Inc Illumina. TruSight One Sequencing Panel Kits Training Support. Illumina Propriety, (January), 2016.

82. Claudia Pommerenke, Robert Geffers, Boyke Bunk, Sonja Eberth, and Hans G Drexler. Enhanced whole exome sequencing by higher DNA insert lengths. BMC Genomics, pages 1–5, 2016.

83. Liucun Zhu, Ying Zhang, Wen Zhang, Sihai Yang, Jian-Qun Chen, and Dacheng Tian. Patterns of exon-intron architecture variation of genes in eukaryotic genomes. BMC Genomics, 10(1):47, 2009.

84. Meena Kishore Sakharkar, Vincent T K Chow, and Pandjassarame Kangueane. Distributions of exons and introns in the human genome. In silico biology, 4(4):387–93, 2004.

85. Mari E K Niemi, Hilary C Martin, Daniel L Rice, Giuseppe Gallone, Scott Gordon, Martin Kelemen, Kerrie McAloney, Jeremy McRae, Elizabeth J Radford, Sui Yu, Jozef Gecz, Nicholas G Martin, Caroline F Wright, David R Fitzpatrick, Helen V Firth, Matthew E Hurles, and Jeffrey C Barrett. Common genetic variants contribute to risk of rare severe neurodevelopmental disorders. Nature, 2018.

86. Joseph H. Nadeau. Modifier genes in mice and humans, 2001.

87. Daniel Greene, Sylvia Richardson, and Ernest Turro. Phenotype Similarity Regression for Identifying the Genetic Determinants of Rare Diseases. American Journal of Human Genetics, 2016.

88. Hilary C. Martin, Wendy D. Jones, James Stephenson, Juliet Handsaker, Giuseppe Gallone, Jeremy F. McRae, Elena Prigmore, Patrick Short, Mari Niemi, Joanna Kaplanis, Elizabeth Radford, Nadia Akawi, Meena Balasubramanian, John Dean, Rachel Horton, Alice Hulbert, Diana S. Johnson, Katie Johnson, Dhavendra Kumar, Sally Ann Lynch, Sarju G. Mehta, Jenny Morton, Michael J. Parker, Miranda Splitt, Peter D. Turnpenny, Pradeep C. Vasudevan, Michael Wright, Caroline F. Wright, David R. FitzPatrick, Helen V. Firth, Matthew E. Hurles, and Jeffrey C. Barrett. Quantifying the contribution of recessive coding variation to developmental disorders. bioRxiv, 2017.

89. Caroline F. Wright, Tomas W. Fitzgerald, Wendy D. Jones, Stephen Clayton, Jeremy F. McRae, Margriet Van Kogelenberg, Daniel A. King, Kirsty Ambridge, Daniel M. Barrett, Tanya Bayzetinova, A. Paul Bevan, Eugene Bragin, Eleni A. Chatzimichali, Susan Gribble, Philip Jones, Netravathi Krishnappa, Laura E. Mason, Ray Miller, Katherine I. Morley, Vijaya Parthiban, Elena Prigmore, Diana Rajan, Alejandro Sifrim, G. Jawahar Swaminathan, Adrian R. Tivey, Anna Middleton, Michael Parker, Nigel P. Carter, Jeffrey C. Barrett, Matthew E. Hurles, David R. Fitzpatrick, and Helen V. Firth. Genetic diagnosis of developmental disorders in the DDD study: A scalable analysis of genome-wide research data. The Lancet, 385(9975):1305–1314, 2015.

90. Yemisi Bokinni. Kabuki syndrome revisited, 2012.

91. N. Bögershausen and B. Wollnik. Unmasking Kabuki syndrome, 2013.

